# Biomechanics and hydraulics of *Opuntia ficus-indica* roots exposed to extreme drought

**DOI:** 10.1101/2022.05.01.490238

**Authors:** Cesar Barrientos-Sanhueza, Vicente Hormazabal-Pavat, Danny Cargnino-Cisternas, Italo F. Cuneo

**Author notes:** Corresponding author: Italo F. Cuneo. Cesar Barrientos-Sanhueza and Vicente Hormazabal-Pavat contributed equally to this manuscript.

## Abstract

Succulent plants possess traits that allow them to complete physiological functions under extreme environments and root are at the frontline of the stress: the drying soil. Previous works in succulent plants have reported the extraordinary reversible mechanism of root shrinkage that disconnects plants from drying soils, reestablishing the hydraulic connection when water availability is restored. Yet, this rectifier-like mechanism would require complex biomechanical and hydraulic control at organ, tissue, and cell level. In here we evaluated the changes in hydraulic and mechanical behavior of *Opuntia* fine roots under extreme drought stress. Using a combination of techniques, we found that fine roots get more elastic as drought stress gets more extreme, allowing cells to modify their shape while preventing permanent damage. Furthermore, we found abrupt decreases in *Lp*_*r*_, that coincided with increased root shrinkage, suberin deposition and structural damage inside the endodermis via lacunae formation and possibly cell wall folding. Our data suggest that, in drought stressed succulent plants, the biomechanics of organs, tissues, and possibly cell walls are deeply coupled with belowground hydraulics, highlighting the need to continue working on deciphering the physiological mechanism that governs the interplay between mechanics and hydraulics at cell level in fine roots during drought.

## 1. INTRODUCTION

Succulent plants possess fascinating anatomical traits (e.g., highly vacuolated parenchyma cells with thin-elastic primary cell walls) that allow them to store water while maintaining high water potentials (Ψ) at tissue level (Gibson and Nobel, 1986; Fradera-Soler et al., 2022). Organ high Ψ is related to the hydraulic capacitance of tissues (*C)*, that is the ability to maintain and release storage water from one tissue to another providing a hydraulic mechanism to avoid the effects of extreme drought (e.g., high values of hydraulic capacitance at turgor loss point, *C*_tlp_) (North et al., 2022; Bartlett et al, 2021). Root systems of succulent plants are in direct contact with stressful microenvironments (i.e., soil-root interface), where the hydraulic resistance can reach up to 90%, generating true bottlenecks in the hydraulic pathway of the SPAC (North and Nobel, 1995; Tsuda and Tyree, 1997; Rodriguez-Dominguez and Brodribb, 2020; Bartlett et al., 2021). Yet, most of the information regarding structural traits that relates to drought resistance in succulent plants has been obtained from the study of aerial organs (Fradera-Soler et al., 2022; Ahl et al., 2019).

Changes in root biomechanical and hydraulic behavior during drought have been linked to anatomical and, specifically, cell wall polysaccharides adjustments (Ahl et al., 2019; North and Nobel, 1991; Loades et al., 2013). One of the most evident changes is the increase in suberization in cell layers outside the stele (e.g., pericycle and cortex; North and Nobel, 1991; Karahara et al., 2004), that increase the hydraulic resistance within roots, negatively affecting water storage (i.e., capacitance) that is necessary to endure under extreme drought periods (Niklas, 1992; Kramer and Boyer, 1995). Root hydraulic conductivity (*Lp*_*r*_) decreases both by the reduction of cell walls permeability caused by suberization, but also by cell wall mechanical failure known as cortical lacunae, hydraulically disconnecting the water flow at the soil-root interface (Zimmermann, 1983; North and Nobel, 1992; Cuneo et al., 2016; Cuneo et al., 2021). Furthermore, low water potential of the vapor phase surrounding the roots creates air gaps at the soil-root interface by decreasing root diameter up to 40% at -10 MPa (i.e., root shrinkage), leading to an even larger reduction of water uptake, water content, and *C*_tlp_ of root cells (North and Nobel, 1992; Nobel and Cui, 1992; Fradera-Soler et al., 2022). Root biomechanical properties are relevant to understand their adaptation to different environments and these properties are influenced by hydrostatic pressure (i.e., cell turgor) and cell wall material composition (Borowska-Wykret and Kwiatkowska, 2018; Ahl et al., 2019; Fradera-Soler et al., 2022). Strength, function, and properties of plant cell walls are linked to their major constituting fundamental blocks, polysaccharides (e.g., cellulose, hemicellulose, and pectins; Ahl et al., 2019; Cosgrove, 2016). An important part of the pectin-rich matrix (i.e., polysaccharides) is located between cells in the apoplastic zone, a local transport route of both water and solutes that could play an important role in how plant tissues respond to drought (Albersheim et al., 2011; Ahl et al., 2019). In a recent original study from Bartlett et al. (2021) it has been highlighted the need to deepen the current knowledge about structural and biochemical adjustments that lead to maintain greater root volume since these traits might have major implication on belowground hydraulics during drought. North (2022) highlighted the intriguing relationship between biochemical/biophysical properties of root cell walls and root capacitance that should be further investigated. In this regard, it has been observed in succulent *Aloe* species that the ability to overcome periodically dry environments is related to regular folding patterns of cell walls (i.e., predetermined cell wall mechanics) that is preceded by biochemical adjustment in the cell wall, allowing its flexibility and preventing damage by drought induced cell shrinkage (Ahl et al., 2019). However, the role of structure and its modification under different periods of extreme drought linked to the root biomechanical properties remain uncertain.

In this study, we hypothesized that exposure to extreme drought will change *Opuntia ficus-indica* roots internal cellular structure by forming lacunae-like damages and suberin deposition in the pericycle and these changes would lead to a modification of the hydraulic and mechanical properties of *Opuntia ficus-indica* roots. The main goal of this study was to evaluate the changes of the hydraulic and mechanical behavior of *Opuntia ficus-indica* young fine roots under extreme drought stress on a 10-, 30-, and 45-days’ time exposure. To test our hypothesis, we used a combination of hydraulic measurements involving a root pressure probe, biomechanical to tests to determine root breaking strain, tensile strength, breaking stress, breaking E, energy strain, and finally using bright field microscopy imaging we analyzed the anatomical and structural changes of Opuntia ficus-indica roots under well-watered and extreme drought treatments. The results of this study provide further knowledge about the relation between structural, mechanical, and hydraulic changes in *Opuntia ficus-indica* roots, thus, improving knowledge on the adaptation of roots to extreme environments.

## 2. MATERIAL AND METHODS

### 2.1 Plant material and establishment

Thirty mature *Opuntia ficus-indica* cladodes averaging 27 ± 3 cm length were collected from Quillota, Valparaíso Region, in Chile. Cladodes were disinfected with ethanol 70% (v/v) and the basal nodes were soaked in 1% (w/v) rooting solution (IAA), placed in a 3.5 L plastic pot filled with growth medium (100% coarse quartz sand) (Supplementary materials, Figure S1A), and maintained for two weeks in a growth chamber until there were signs of root initiation and proliferation (Supplementary materials, Figure S1B) (North & Nobel, 1991; Cuneo et al., 2016). After the acclimatation time (i.e., two weeks), 18 plants were selected and cleaned. The field capacity of the growth medium was determined by the wet and dry weight of the coarse quartz sand, which was saturated with filtered water and dried at 105ºC in an oven for 24 hours. (Barrientos-Sanhueza et al., 2022). According to the wet and dry mass difference the field capacity was equal to 195 ml kg^-1^ of growth medium. Plants were irrigated to field capacity each 3 days.

### 2.2 Plant growth and treatment setup

All the experiments were carried out in the Plant and Soil Biophysics and Biomechanics lab at Pontificia Universidad Católica de Valparaíso, Chile. Three levels of drought were stablished: 10, 30 and 45 days. Nine Opuntia cladodes with triplicates for well-watered (i.e., control) and drought treatments were selected (26.17 cm ± 3.15 cm length and 27.55 cm ± 2.04 cm length, respectively). Plant growth was maintained under environmental controlled conditions (approximately 16ºC to 28ºC temperature, 35% relative humidity with natural light 500-1100 µmol photons m^-2^ s^-1^ PAR, supplemented light 400-600 µmol photons m^-2^ s^-1^ PAR was provided with 600-W metal-halide bulbs (PIRANHA, China), and 16-h-light/8-h-dark cycle) and were irrigated with filtered water three times per week for 45 days. Control (C) and drought (D) measurement treatments were carried out at 10, 30 and 45 days (C10, C30, C45 and D10, D30, D45, respectively). An external heating mat with a temperature of 30°C was put under the drought treatments plastic pots after a week for 24 hours to accelerate the evaporation of water from the coarse quartz sand. After 10, 30 and 45 days the pots were opened, and the root area was washed with filtered water. Roots were carefully excised under water, cleaned, and then storage on demineralized water. Then, roots of 80 to 110 mm length were selected for mechanical, hydraulic, and microscopy experiments. (Nobel & Cui, 1992; North & Nobel, 1991).

### 2.3 Root tensile strength

Tensile strength measurements (roots n = 3) were carried out using a Texture analyzer (Model Ta.XT *plusC*, Stable Micro Systems Ltd., Surrey, UK) equipped with mini tensile grips (A/MTG, Stable Micro Systems Ltd., Surrey, UK). Uniaxial tensile strength was applied to stretch until failure test approach (Bidhenhi and Geitmann, 2018). Sampled roots were selected, and each root end were attached to the mini tensile grips using Parafilm® M film (Bemis Company Inc., USA), then inserted into 200 µl micropipette tips and glued with superglue (La Gotita®, FENEDUR S.A., Uruguay) to improve grip. The texture analyzer software was set as follows: pre-test = 0.01 mm s^-1^, speed-test = 0.2 mm s^-1^, post-test = 0.05 mm s^-1^, force = 0.05 N (Barrientos-Sanhueza et al., 2021; Barrientos-Sanhueza et al., 2022). Trigger force were set in force, and the tensile force was recorded in Newton (N) at a yield force of 5 N to record with accuracy the mechanical behavior of treated roots (Barrientos-Sanhueza et al., 2022). Besides, time and distance were recorded too. Data were considered only when the break took place in the middle part of the root (Supplementary material S3). Finally, data were then transformed into Mega Pascals (MPa) and axial strain. This data was used to the stress-strain graphs and to estimate breaking strain, tensile strength (N), breaking stress (MPa), and Young’s module (E, MPa).

To calculate the later, we estimated the engineering stress-strain data, as follows:

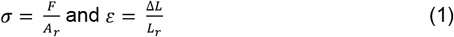

where *σ* is engineering stress (MPa) and *ε* is the engineering strain. *A_r_* is the root cross-section area (m^2^) and *L_r_* is the original length of the root (m). Δ*L* = L-L is the uniaxial displacement of the root (Bidhendi et al., 2019; Niklas, 1992). Finally, to calculate the amount of energy, that must be introduced into the system (i.e., plant root) to modify the biophysical properties of cell walls from cell to organ level, the strain energy (U) was calculated as follow (Niklas, 1992; Barrientos-Sanhueza et al., 2022):

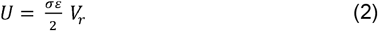

where *σ* is stress (MPa), *ε* is strain, and *V_r_* is root volume (m). Besides, U can be directly calculated for linear elastic materials. Nevertheless, for non-elastic and plastic materials, U must be estimated from the area under the stress-strain diagram (i.e., area under the curve) (Barrientos-Sanhueza et al., 2022).

### 2.4 Root pressure (*P*_*r*_) and hydraulic conductivity (*Lp*_*r*_) of *Opuntia* fine roots

To measure ***P***_***r***_ and ***Lp***_***r***_ (i.e., water movement across the root), a root pressure probe system was used according to the procedure described by Knipfer et al. (2007). First, root segments were tightly connected to the root pressure probe system inserting them through a cylindrical hydrophilic dental impression silicon seal (made from liquid silicone material; Speedex kit, Coltene, Germany) surrounded by a stainless-steel clamp and the seal was tightened softly with a screwdriver until there was a small pressure increase (North and Nobel, 1991). The seal was connected to a microfluidic system to optimize data accuracy and collection. In brief, a pressure sensor of 1035 mbar (Pressure Sensor, MPS2, Elveflow, France) connected to a sensor reader (Sensor Acquisition Module, MFS2, Elveflow, France) was coupled to a microfluidic interface and to the silicon seal. Readings from the sensor were captured and saved with the Elveflow ESI software version 3.02.05. Steady-state root pressures were reached and recorded before continuing with the measurements, and the half-times of the final relaxations were at least 10-fold smaller than the initial relaxations (Peterson et al., 1993). To estimate the compressibility of the measurement system, which is given by the system’s modulus of elasticity (β), it was necessary to perform several volume changes (Δ*V_s_*), which resulted in changes in root pressure (ΔP_r_) (Knipfer et al., 2007; Steudle et al., 1987). The system’s elastic modulus (β) is a very important factor for root pressure probe accuracy measurements. The lower the absolute value or higher the relation of 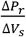 the higher the sensitivity by which *Lp*_*r*_ can be measured (Steudle et al., 1987). The hydraulic conductivity of the root *Lp*_*r*_ of excised *Opuntia* fine roots was evaluated through the half-times (*T*_*1/2*_) of hydrostatic pressure relaxation (Knipfer et al., 2007; Steudle et al., 1987; Peterson et al., 1993). In the hydrostatic experiments, the root was bathed in filtered water and, with a syringe pump (Longer Pump, LSP01-3A, Longer Precision Pump Co., Ltd., China), several pressure pulses were applied, increasing, or decreasing the steady-state root pressure, maintaining the system’s volume constant until the equilibrium of the pressure was restored. *Lp*_*r*_ was estimated as fallows (Steudle and Peterson, 1998; Knipfer et al., 2007; Steudle et al., 1987):

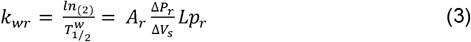

where *k_wr_* is the rate constant of water interchange through the root; 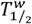 is the half-time of hydrostatic pressure relaxation; 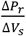 is the elastic coefficient of the measuring system (in MPa m^-3^); and A is the effective surface area of the root (in m^-2^). To estimate *A_r_*, the effective surface area (m^2^) of the different root segments was measured using *Fiji* imaging-processing software (www.fiji.sc, ImageJ) (Cuneo et al., 2018; Supplementary material S2).

### 2.5 Bright field microscopy imaging

Root anatomy was observed on free-hand cross-sections that were made from the same root developmental regions (i.e., “Transition zone”, 5-6 cm from root tip) using fresh razor blades (Cuneo et al., 2016). For general root anatomy identification of lignified cell walls or polysaccharides rich in carboxy groups, root sections were stained with 0.01% (w/v) toluidine blue O for 5 min and view under bright light (O‘Brien et al., 1964; Knipfer and Fricke, 2010; Barrios-Masias et al., 2015; Gambetta et al., 2013; North and Nobel, 1991). To identify suberin and lipid deposits (i.e., red color), the root sections were stained with Sudan Red 7B for 2 h. A solution of Sudan Red 7B at 0.01% with polyethylene glycol (PEG 400): glycerol (1:1; v/v) was prepared (Brundrett et al., 1991; Knipfer and Fricke, 2010; Li et al., 2007). Toluidine blue O and Sudan Red 7B stained sections were mounted on slides with diH_2_O and glycerol under the microscope, respectively (Cuneo et al., 2016; Cuneo et al., 2020). Finally, stained mounted root sections images were observed and captured with a Leica MC170HD digital camera attached to a Leica DMIL LED inverted microscope (Leica Microsystems, Wetzlar, Germany) (Barrientos-Sanhueza et al., 2022; Barrientos-Sanhueza et al., 2021).

### 2.6 Statistical analysis

Analysis of variance (ANOVA) and *t-tests* were performed using R version 4.2.0 statistical computing environment (R Core Team, 2022) with the aid of the CAR software package (Fox & Weisberg, 2011). The Shapiro–Wilk and Levene tests were used to check for normality and homogeneity of variance, respectively. Tukey’ honest significant difference test was used to determine significant differences among treatments.

## 3. RESULTS

### 3.1 Tensile strength test

Root diameter varied among *Opuntia* treated roots from 4.6 mm to 7.8 mm. There was no statistical difference between the C10-D10 (6.90 × 10^−3^ ± 4.51 × 10^−4^ and 6.50 × 10^−3^ ± 5.13 × 10^−4^, respectively) and C30-D30 (5.90 × 10^−3^ ± 8.14 × 10^−4^ and 5.30 × 10^−3^ ± 1.73 × 10^−4^, respectively) in root diameter. However, statistical differences were found between C45-D45 root diameter (7.37 × 10^−3^ ± 7.31 × 10^−4^ and 5.77 × 10^−3^ ± 9.28 × 10^−4^, respectively). Root shrinkage progressively increased over a 10-, 30-, and 45-days period (6%, 10%, and 22%, respectively) (Figure 1A, B, and C). We used the tensile strength test to quantify the mechanical behavior of the different treated *Opuntia* roots on the stress-strain relationship, breaking strain, tensile strength (N), breaking stress (MPa), and Young’s module as breaking E (MPa) there was significant differences between the treatments (P≤0.05, Student‘s *t*-test) (Table 1). Typical elastic-plastic behavior was observed in the stress-strain curves, with steeped initial curve before the plastic zone after the yield point (Figure 2). During the mechanical measurements, several snapping sounds were identified, revealing the cortex break and ultimate tensile failure (Figure 2). Tensile strength was calculated on tensile load over the sample, and breaking stress was calculated on root cross-sectional area, while breaking E was calculated by the stress-strain relationship which is part of the linear portion. Breaking strain had little variation among treatments. C10 was almost 1-fold higher compared with D10 (3.22 × 10^−2^ ± 5.40 × 10^−3^ SE and 2.40 × 10^−2^ ± 1.08 × 10^−3^ SE, respectively), but the rest of the treatments displayed similar values and no significant differences were found (P≤0.05). Significant differences were found in tensile strength, breaking stress, and breaking E measurements (*p* < 0.01). C45 (3.47 ± 1.33 N SE) displayed almost 3-fold higher tensile strength compared with D45 (1.08 ± 0.25 N SE), and C10 (1.28 ± 0.14 N SE) showed almost 1-fold higher tensile strength compared with D10 (0.52 ± 0.10 N SE). However, D30 displayed 0.4 N higher value compared with C30 (1.35 ± 0.39 N SE and 0.95 ± 0.45 N SE, respectively). Breaking stress values of C45 (1.50 × 10^−2^ ± 4.67 × 10^−3^ MPa SE) were 2.3-fold higher than D45 (6.36 × 10^−3^ ± 2.06 × 10^−3^ MPa SE). C10 and D10 displayed the same behavior (7.55 × 10^−3^ ± 1.15 × 10^−3^ MPa SE and 3.23 × 10^−3^ ± 4.37 × 10^−4^ MPa SE, respectively). Conversely, D30 showed 2-fold higher breaking stress compared with C30 (9.73 × 10^−3^ ± 1.78 × 10^−3^ MPa SE and 4.84 × 10^−3^ ± 2.04 × 10^−3^ MPa SE, respectively). To classify and identify the capacity of a body to absorb and subsequently release strain energy before it breaks (i.e., elastic resilience before failure), strain energy (U) was estimated. At 10-days, well-watered (i.e., C10) treatment showed higher strain energy, reaching almost 4-fold (1.88 × 10^−9^ ± 5.92 × 10^−10^ J SE) compared with extreme drought treatment (i.e., D10) (4.75 × 10^−10^ ± 2.50 × 10^−10^ J SE) (Figure 2A, B). Nevertheless, there were big differences between the stress and strain behaviors. D10 displayed higher stress value (0.0020 MPa) compared with C10 (0.00064 MPa), but lower strain values (0.012 and 0.023, respectively). At 30-days, no changes were observed on stress behavior between well-watered and extreme drought treatments (1.40 × 10^−9^ ± 6.71 × 10^−10^ J SE and 2.78 × 10^−9^ ± 1.14 × 10^−9^ J SE), but D30 still had a larger strain behavior compared with C30 (0.032 and 0.027, respectively). Another scenario happened at 45-days, where D10 displayed 2-fold higher strain energy (1.81 × 10^−10^ ± 3.18 × 10^−11^ J SE) compared with C10 (3.91 × 10^−10^ ± 1.47 × 10^−10^ J SE). However, C45 showed larger stress value compared with D45 (0.0014 MPa and 0.00059 MPa, respectively). Finally, to estimate the elastic properties of ultimate tensile failure of different treated roots, Young’s module as breaking E (MPa) was analyzed, and we found statistical differences (P≤0.05, Student‘s *t*-test) at 10 days. C10 showed almost 2-fold higher than D10 (2.36 × 10^−2^ ± 4.71 × 10^−4^ MPa SE and 1.35 × 10^−2^ ± 1.89 × 10^−3^ MPa SE, respectively). Besides, C45 displayed the higher value of elasticity at failure compared with D45 (7.00 × 10^−2^ ± 2.50 × 10^−2^ MPa SE and 3.19 × 10^−2^ ± 1.22 × 10^−2^ MPa SE, respectively). Nevertheless, at 30 days the relation was reversed. C30 showed 1.4-fold higher value than C30 (2.44 × 10^−2^ ± 2.82 × 10^−3^ MPa SE and 1.73 × 10^−2^ ± 6.72 × 10^−3^ MPa SE, respectively), the same behavior previously reported in tensile strength and breaking stress (Table 1).

**Figure 1.**
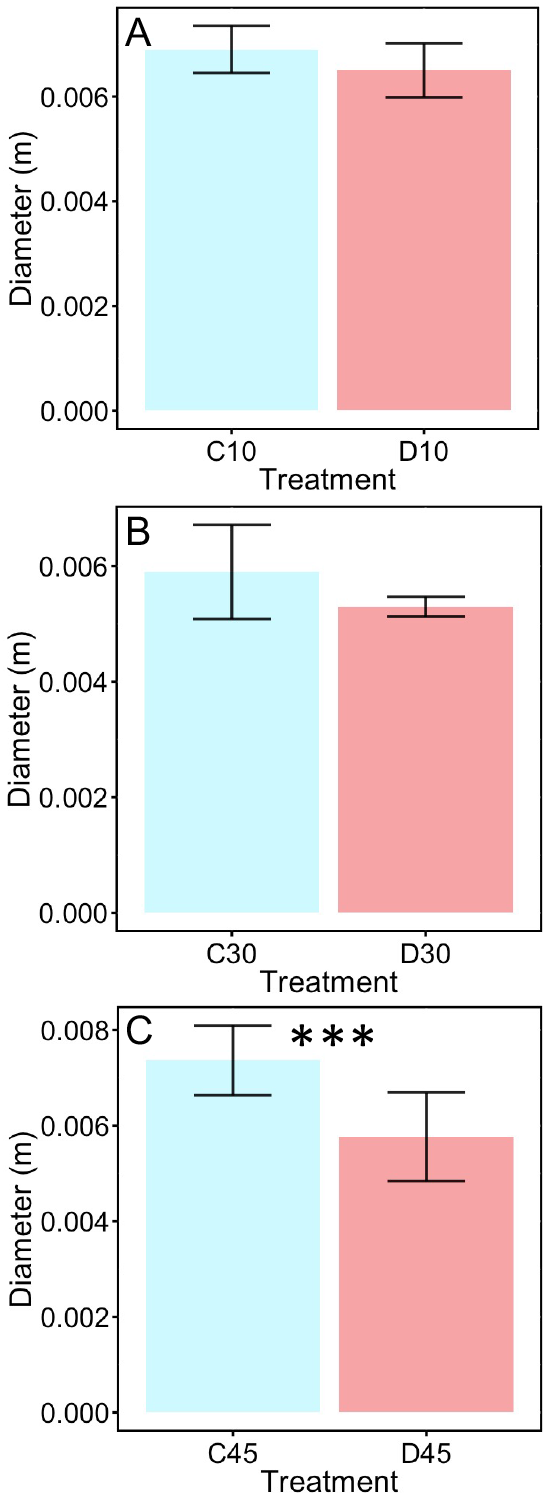
*Opuntia* root diameter of the different treatments. (A), *Opuntia* roots measured at 10 days. (B), *Opuntia* roots measured at 30 days. (C), *Opuntia* roots measured at 45 days. Blue and red colour indicate well irrigated and extreme drought treatments, respectively. Statistical differences were determined with *t*-test. *p* < 0.01 = **; p < 0.001 = *** (n=3).

**Table 1.**
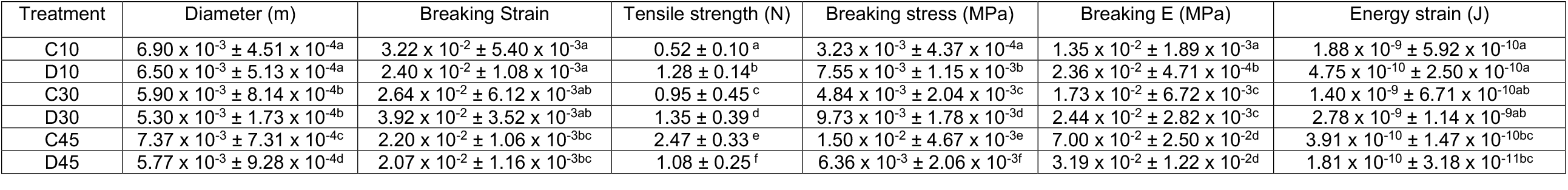
Biomechanical behavior of roots under well-watered and extreme drought period treatments. Data are means ±SE (n=3). Similar letters denote nonsignificant differences. Statistical difference is indicated by different superscripts (P≤0.05, Student‘s *t*-test).

| Treatment | Diameter (m) | Breaking Strain | Tensile strength (N) | Breaking stress (MPa) | Breaking E (MPa) | Energy strain (J) |
| --- | --- | --- | --- | --- | --- | --- |
| C10 | $6.90 \times 10^{-3} \pm 4.51 \times 10^{-4a}$ | $3.22 \times 10^{-2} \pm 5.40 \times 10^{-3a}$ | $0.52 \pm 0.10^a$ | $3.23 \times 10^{-3} \pm 4.37 \times 10^{-4a}$ | $1.35 \times 10^{-2} \pm 1.89 \times 10^{-3a}$ | $1.88 \times 10^{-9} \pm 5.92 \times 10^{-10a}$ |
| D10 | $6.50 \times 10^{-3} \pm 5.13 \times 10^{-4a}$ | $2.40 \times 10^{-2} \pm 1.08 \times 10^{-3a}$ | $1.28 \pm 0.14^b$ | $7.55 \times 10^{-3} \pm 1.15 \times 10^{-3b}$ | $2.36 \times 10^{-2} \pm 4.71 \times 10^{-4b}$ | $4.75 \times 10^{-10} \pm 2.50 \times 10^{-10a}$ |
| C30 | $5.90 \times 10^{-3} \pm 8.14 \times 10^{-4b}$ | $2.64 \times 10^{-2} \pm 6.12 \times 10^{-3ab}$ | $0.95 \pm 0.45^c$ | $4.84 \times 10^{-3} \pm 2.04 \times 10^{-3c}$ | $1.73 \times 10^{-2} \pm 6.72 \times 10^{-3c}$ | $1.40 \times 10^{-9} \pm 6.71 \times 10^{-10ab}$ |
| D30 | $5.30 \times 10^{-3} \pm 1.73 \times 10^{-4b}$ | $3.92 \times 10^{-2} \pm 3.52 \times 10^{-3ab}$ | $1.35 \pm 0.39^d$ | $9.73 \times 10^{-3} \pm 1.78 \times 10^{-3d}$ | $2.44 \times 10^{-2} \pm 2.82 \times 10^{-3c}$ | $2.78 \times 10^{-9} \pm 1.14 \times 10^{-9ab}$ |
| C45 | $7.37 \times 10^{-3} \pm 7.31 \times 10^{-4c}$ | $2.20 \times 10^{-2} \pm 1.06 \times 10^{-3bc}$ | $2.47 \pm 0.33^e$ | $1.50 \times 10^{-2} \pm 4.67 \times 10^{-3e}$ | $7.00 \times 10^{-2} \pm 2.50 \times 10^{-2d}$ | $3.91 \times 10^{-10} \pm 1.47 \times 10^{-10bc}$ |
| D45 | $5.77 \times 10^{-3} \pm 9.28 \times 10^{-4d}$ | $2.07 \times 10^{-2} \pm 1.16 \times 10^{-3bc}$ | $1.08 \pm 0.25^f$ | $6.36 \times 10^{-3} \pm 2.06 \times 10^{-3f}$ | $3.19 \times 10^{-2} \pm 1.22 \times 10^{-2d}$ | $1.81 \times 10^{-10} \pm 3.18 \times 10^{-11bc}$ |

**Figure 2.**
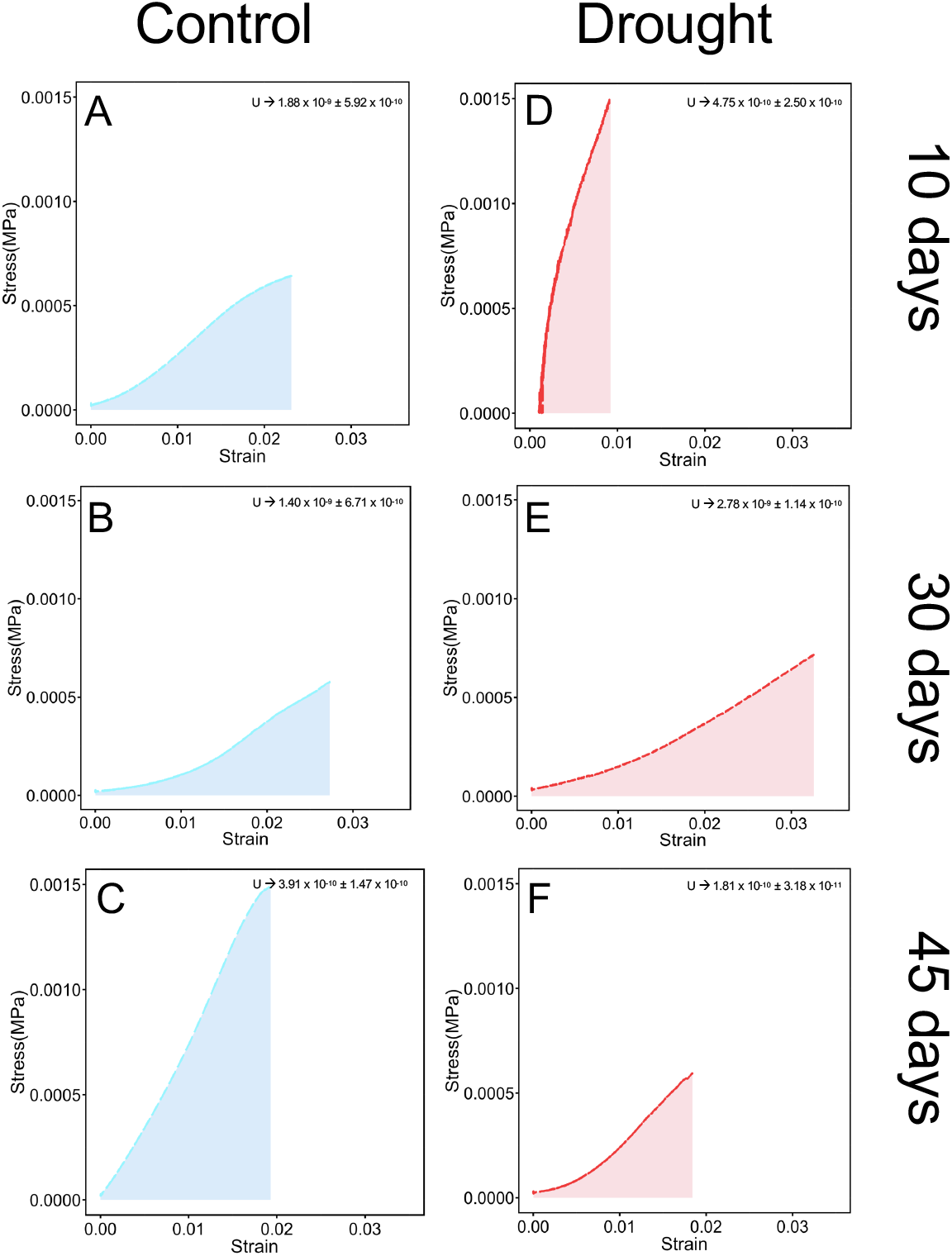
Tensile strength curves representing the mechanical properties of *Opuntia* roots. Figure shows the effect of different drought treatments over the stress-strain (s-s) behavior and the area under the curve that represents the strain energy (U). (A-F) Strain energy values are in the upper right section. (A-C) Blue lines and area represent well-watered treatments at 10-, 30-, and 45-days. (D-F) Red lines and area represent extreme drought treatment at 10-, 30-, and 45-days. Data are means ±SE (n=3).

### 3.2 Hydraulic conductivity test

Significant differences of *Lp*_*r*_ were found among 10-, 30-, 45-day between well-watered and extreme drought treatments (p = 0.00185 and p = 8.67 × 10^−5^, respectively) (Figure 3). We observed a reduction of 48% in *Lp*_*r*_ between 10- and 30-day in well-watered plants (2.27 × 10^−8^ ± 3.55 × 10^−9^ m s^-1^ MPa^-1^ and 1.18 × 10^−8^ ± 3.39 × 10^−9^ m s^-1^ MPa^-1^, respectively) while no differences in *Lp*_*r*_ were found between 30- and 45-day in well-watered plants (1.18 × 10^−8^ ± 3.39 × 10^−9^ m s^-1^ MPa^-1^ and 1.17 × 10^−8^ ± 1.67 × 10^−9^ m s^-1^ MPa^-1^, respectively). Besides, a reduction of 67% and 54% in *Lp*_*r*_ between 10/30-day (9.68 × 10^−9^ ± 1.62 × 10^−9^ m s^-1^ MPa^-1^ and 3.53 × 10^−9^ ± 8.64 × 10^−10^ m s^-1^ MPa^-1^, respectively) and 30/45-day (3.53 × 10^−9^ ± 8.64 × 10^−10^ m s^-1^ MPa^-1^ and 1.49 × 10^−9^ ± 2.47 × 10^−10^ m s^-1^ MPa^-1^, respectively) in extreme the drought treatment was found, respectively. During the experiments, 10-days of drought was enough to observe a pronounced reduction in *Lp*_*r*_ (i.e., 57% of reduction compared with well-watered plants), while the *Lp*_*r*_ of 30- and 45-days extreme drought treatments decreased, even more, to 73% and 87%, respectively, compared with well-watered plants.

**Figure 3.**
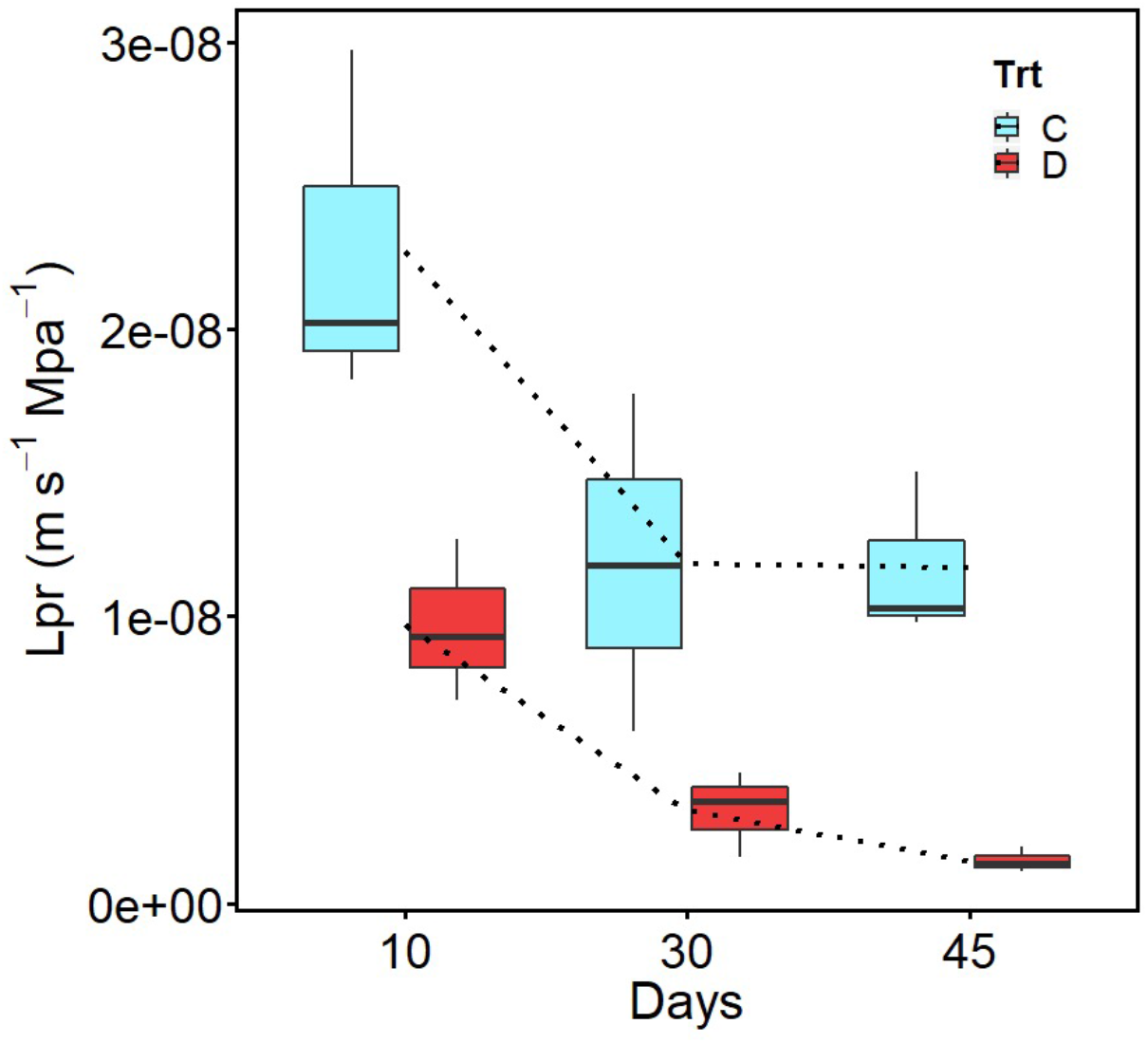
Extreme drought-induced changes in *Opuntia* radial root hydraulic conductivity (Lp_r_). Blue and red color indicate well irrigated and extreme drought treatments at 10-, 30-, and 45-days, respectively. Data are means ±SE (n=3).

### 3.3 Microscope imaging

Bright field microscopy was used to visualize the different root cellular composition and the materials that compose them under well-watered and extreme drought conditions. Transition zone (i.e., 5-6 cm from root tip) root images of well-watered plants at 10-days presented a normal inner cell conformation for a 1-month-old *Opuntia ficus-indica* root. Using toluidine blue O, several cell compartments were identified. Epidermis (i.e., external tissue) was composed of compact cells, and some cells producing root hairs. Immediately below the epidermis we observed the hypodermis, which is the outermost cortical cell layer. Then continues the layer of cells of the cortex and, finally, endodermis (i.e., the innermost cortical cell layer). Inside the endodermis, we observed the vascular cylinder, a complex tissue formed by different materials (e.g., suberin, lignin, polysaccharides, between others). Besides, this cylinder comprises from the outer to the inner zone, a three-cell-layered pericycle, xylem, phloem, and the vascular parenchyma with a polyarchy root vasculature structure with five to eight xylem poles, a classic characteristic of platyopuntia (Freeman, 1969; Nobel, 2002; Figure 4).

**Figure 4.**
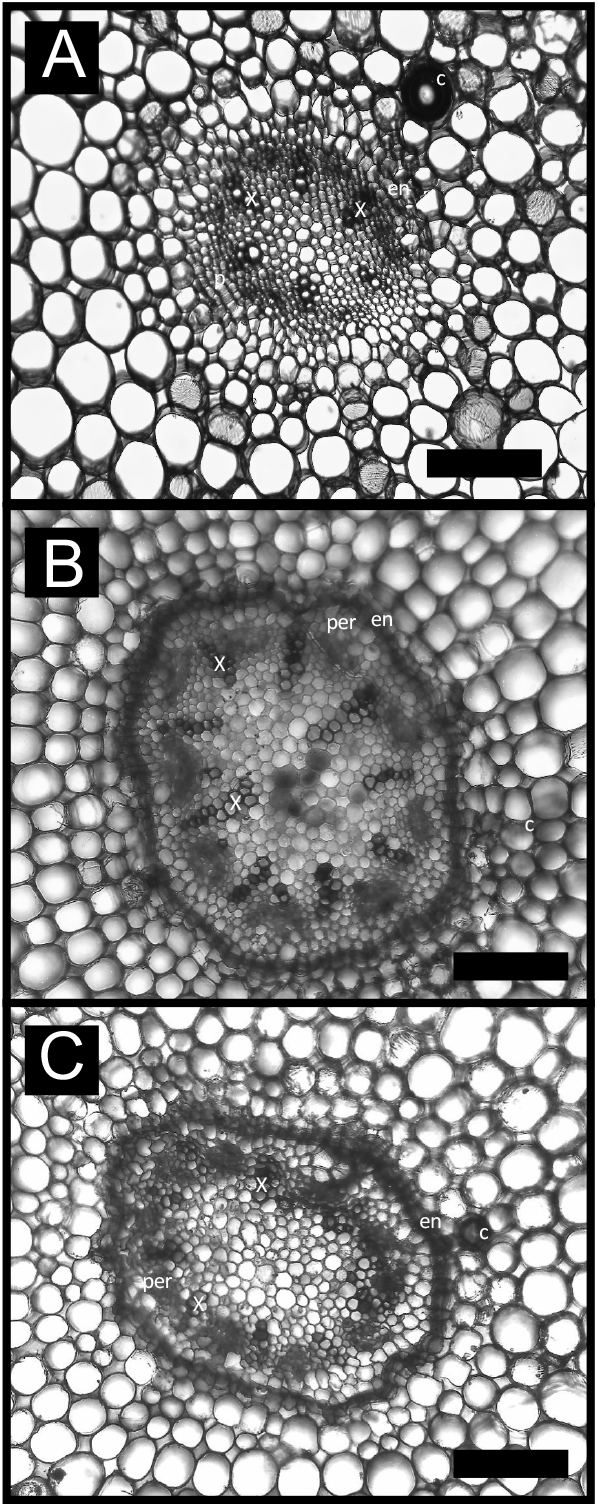
Free-hand cross-section of well irrigated treatments of *Opuntia ficus-indica* roots. (A), shows the central zone of a 10-day-old root. (B), shows the central zone of a 30-day-old root. (C), show the central zone of a 45-day-old root. Samples were stained with toluidine blue O (TBO; in deionized water) and Sudan Red 7B. Gray filters were applied on root images to improve structural and anatomical contrasts. c, Cortex; x, Xylem; en, Endodermis; per, Peridermis. Bars = 100 µm.

Cortical and pericycle cells at 45 days of drying started to fold and losing roundness caused probably by the loss of water (Figure 5B, 6B). Suberin was observed with an intense turquoise color in the radial zone of the endodermis, indicating the casparian strip (Esau, 1965). Also, suberin was present in the radial wall of mature peridermal cells. At 10-days, extreme drought treatment showed the presence of several structures suberized, and the first cell wall folding phenomena was observed (Figure 6A). On the other hand, at 45-days extreme drought treatment, the pericycle and endodermis cortex showed a pronounced cell wall folding phenomenon (Figure 6B). Finally, 45-days of extreme drought treatment caused the failure and plasmolysis of pericycle cell layers close to the casparian strip giving place to pericycle lacunae to form (Figure 5B, 6B), possible due to the loss of cell water content and, consequently, cell roundness.

**Figure 5.**
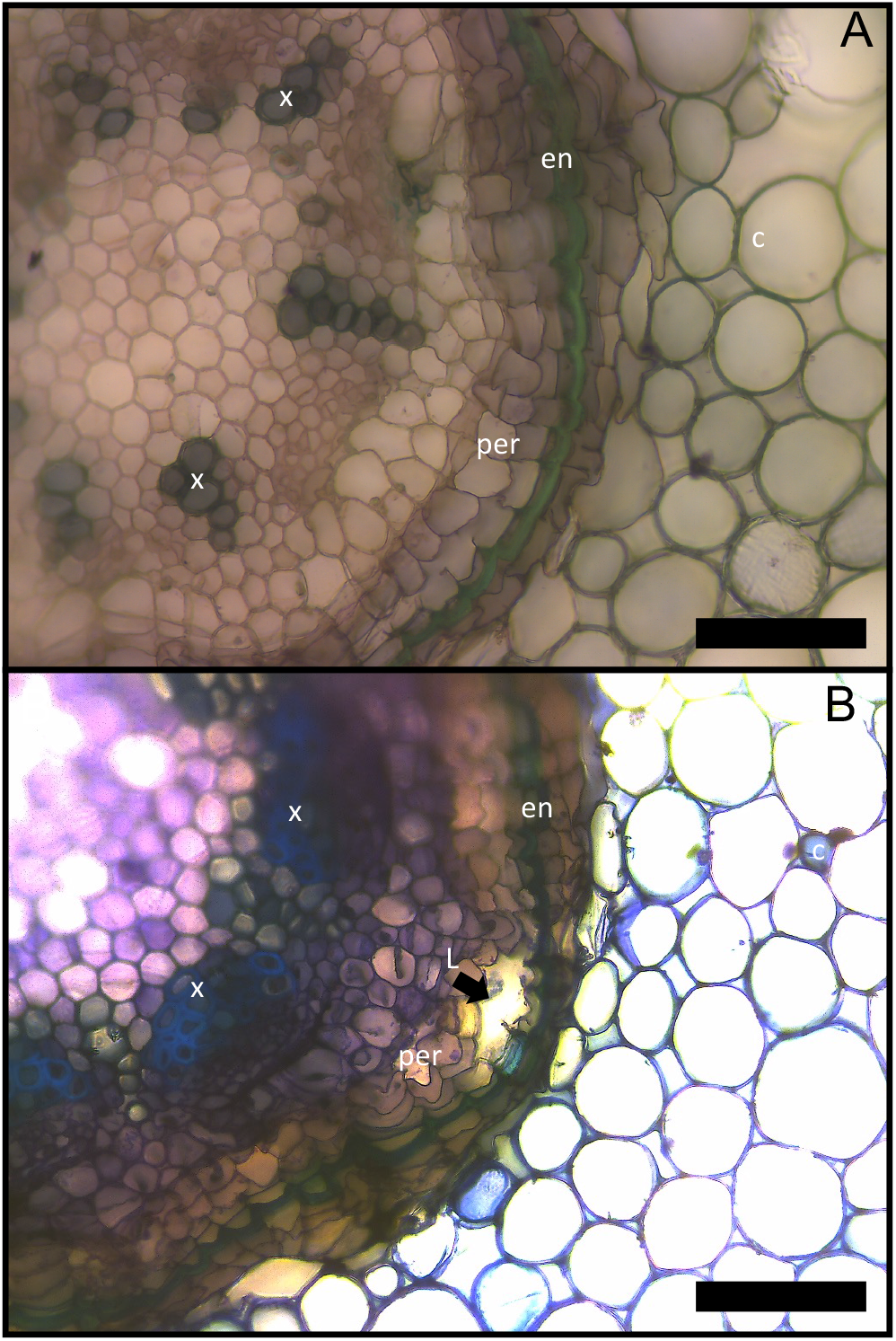
Root cell Wall folding and structural differences of *Opuntia ficus-indica* roots free-hand cross-section. (A), 45-day-old root under well irrigated treatment, without cell damage. (B), 45-day-old root under extreme drought treatment, with cell damage. Samples were stained with toluidine blue O (TBO; in deionized water) and Sudan Red 7B. Black arrow indicates mechanical failure of pericycle, disconnecting part of the endodermis with the stele. c, Cortex; x, Xylem; en, Endodermis; per, Peridermis; L, Lacunae. Bars = 200 µm.

**Figure 6.**
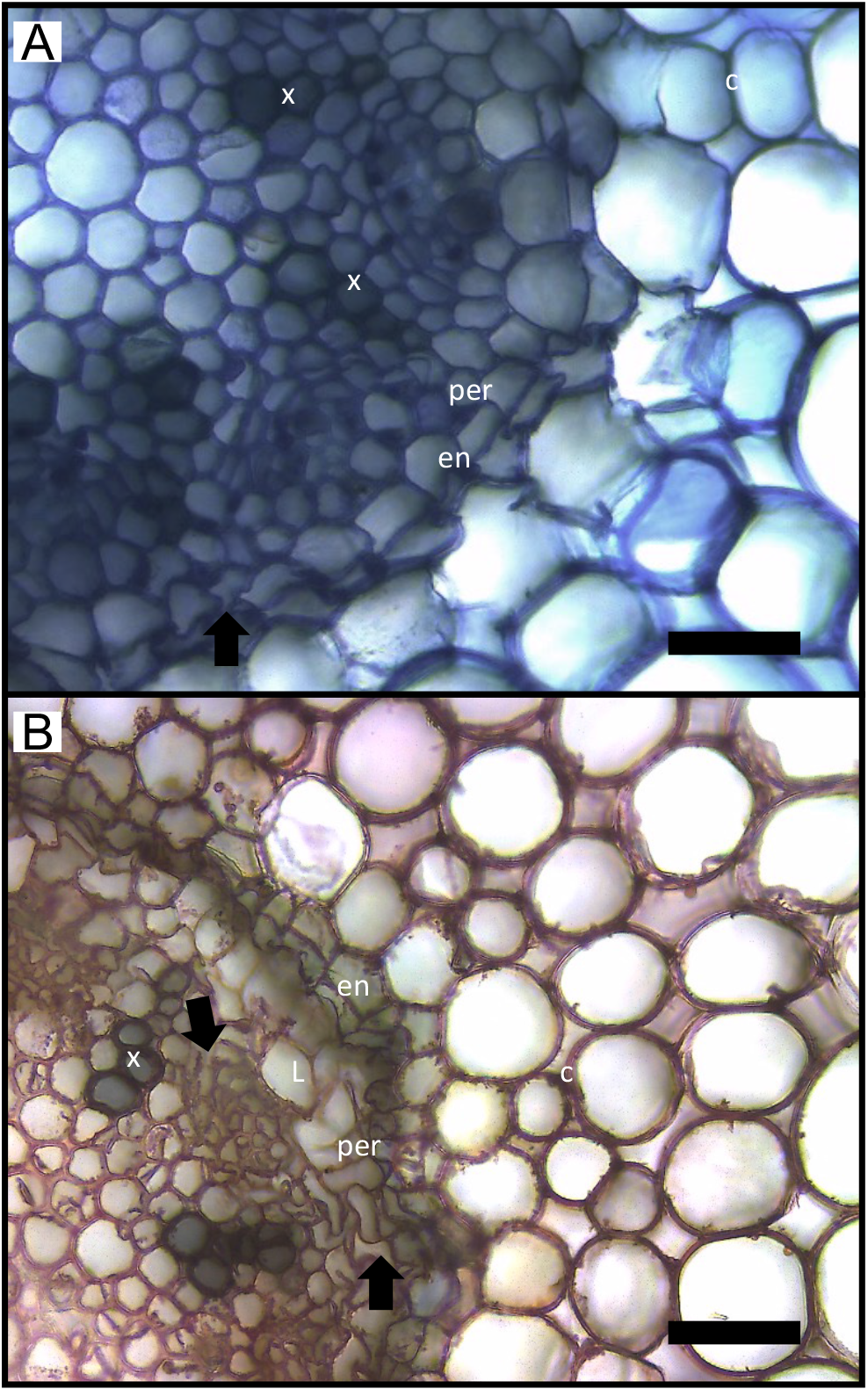
Comparation of free-hand cross-section of *Opuntia ficus-indica* roots at different extreme drought periods. (A), 10-days-old root under extreme drought treatment (D10) stained with toluidine blue O (TBO). (B), 45-days-old root under extreme drought treatment (D45) stained with Sudan red 7B. Black arrows indicate the cell wall folding formation and the presence of cell discontinuity. c, Cortex; x, Xylem; en, Endodermis; per, Peridermis; L, Lacunae. Bars = 100 µm.

## 4. DISCUSSION

Our results provide evidence from hydraulic, mechanical, and anatomical analysis that prolonged extreme drought periods modify the biomechanical and hydraulic behavior of *Opuntia ficus-indica* fine roots. Previous works in succulent plants have reported the extraordinary reversible mechanism of root shrinkage that disconnects plants from drying soils, reestablishing the hydraulic connection when water availability is restored (Nobel and Cui, 1992; North and Nobel, 1997) and our data confirms these findings (i.e., 22% of root shrinkage at 45-days of extreme drought). To perform this rectifier-like behavior, our results highlight that the biomechanics of whole tissues and organs plays a key role, something that has been observed in the past in aerial organs of succulents (Fradera-Soler et al., 2022). We found that fine roots get more elastic as drought stress gets more extreme, a mechanism that might be related to changes in cell wall composition, allowing cells to change their shape while preventing permanent damage. Furthermore, we found abrupt decreases in *Lp*_*r*_, that coincided with observations of suberin deposition and structural damage inside the endodermis via lacunae formation and possibly by cell wall folding modification. Our data suggest that, in drought stressed succulent plants, the biomechanics of organs, tissues, and possibly cell walls are deeply coupled with belowground hydraulics (Bartlett et al., 2021; North, 2022), highlighting the need to continue working on trying to decipher the physiological mechanism that governs the interplay between mechanics and hydraulics at cell level in fine roots during drought.

Previous work on succulent tissues have reported the effects of drought over osmotic adjustment and cell shrinkage and it‘s relation to cell wall mechanics and water uptake (Westermaies, 1884; Haberlandt, 1904, Engmann, 1934; North and Nobel, 1991; North and Nobel, 1992). *Opuntia ficus-indica* root cell walls are characterized by having thin and highly flexible primary cell walls, especially in cortex tissues (Figure 4; North and Nobel, 1992; Fradera-Soler et al., 2022). On the other hand, succulent roots need support tissues such as fibers, suberin, and lignin from secondary growth to support the high hydrostatic pressure generated by high cell turgor (Niklas, 1992; Mauseth, 2006; Bobich and North, 2009). Besides, the nature of succulent plants is related to maintain a relatively high-water potential (Ψ) at organ and cell level during extended drought periods (Nobel and Jordan, 1983; North and Nobel, 1992). Succulent ability to maintain high Ψ values can be understood in terms of hydraulic capacitance (*C)*, specially, high values of hydraulic capacitance at turgor loss point (*C*_tlp_) allows to maintain and release storage water from one organ or tissue to another, and consequently, could support and avoid water stress (North, 2022; Ahl et al., 2019). Our results suggest that, under extreme drought events, the biophysical properties of cell walls are modified at cell and organ level, generating a series of root biomechanical changes. Tensile strength tests showed that the stress-strain relationship, which is a useful way to quantify the plant material resistance against deformation (Bidhendi and Geitmann, 2018; Bidhendi and Geitmann, 2019), had a tensile strength value 2.4-fold bigger at 10-days drought treatment (i.e., D10) than well-watered condition (i.e., control; C10), same behavior occurs with breaking stress and breaking E values, meaning that D10 improves root strength. However, in terms of energy strain (U), C10 was almost 4-fold bigger than D10 before breaking (Figure 2A, D). Tensile loads applied to the different treated roots add energy to the system, and for elastic-plastic materials this energy produce molecular rearrangement, and in case of strain hardening materials (e.g., plant roots), U aligns the polymer chains (e.g., cell wall polysaccharides) in the direction of externally applied forces until failure (Niklas, 1992). For drought-avoiding succulents (e.g., *Opuntia ficus-indica)*, rapid adaptation is essential for survivance, specially under long drought periods (Fradera-Soler et al., 2022). Rapid adaptation consists of elastic adjustment by decreasing the volumetric modulus of elasticity (ϵ) at cell level, and this elastic adjustment could be involved on the rapid changes of the cell wall polysaccharide structure and geometry (i.e., remodeling), specially of the pectin section (Schulte, 1992; Peaucelle et al., 2011; Bethke et al., 2016; Ahl et al., 2019; Fradera-Soler et al., 2022). The modification of pectin section could be linked to the cell wall folding phenomena that occurs in dehydrated succulent tissues and its consequent effects on hydraulic conductance and conductivity, something that has been largely reported on aerial tissues (e.g., leaves, and shoots) (Ahl et al., 2019; Schmidt and Kaiser, 1987; Nobel, 2006; Sack and Scoffoni, 2013; Brodribb et al., 2007; Gibson and Nobel, 1986; Smith and Nobel, 1986; Buckley, 2015; Buckley et al., 2015). However, little is known about what happened in roots, maybe because roots are much less accessible than shoots (Steudle, 2000).

Water follows an intricated pathway in the soil-plant-atmosphere continuum (i.e., SPAC) (Steudle, 2000). Water pass through the apoplastic and cell to cell pathway in a combined fashion and downward water potentials (Steudle, 2000). Our results confirm previous results indicating that these pathways are modified and interrupted during drought by lignified and suberized cell walls, showing thicker-walled cells (Figure 5 and 6), and coinciding with the abrupted reduction in *Lp*_*r*_ (North and Nobel, 1992). In our work, *Lp*_*r*_ of D45 had a reduction of 87% compared with C45. However, the apparently negative abrupt reduction of *Lp*_*r*_ specially under extreme drought events, could lead to a positive effect on root biomechanics. First, this reduction of root hydraulic conductivity limits the water uptake that is necessary to maintain the water potential at the turgor loss point (*TLP_ψ_*) in succulent tissues generating a change from the regular, spherical, and turgid cell characteristic to a convoluted folding of collapsible cell walls but maintaining a high turgor that allows to preserve the cell membrane-cell wall continuum that prevents the irreversible damage due to several mechanical stresses (see Figure 5B, 6A, B; Fradera-Soler et al., 2022). The change in cell volume, and shape under drought modifies the elastic behavior at cell and organ level, and it is affected by the fibrillar network of microfibrils within the cell wall matrix under different loads, and in our case, of different hydrostatic pressures depending on how full the matrix is hydrated (Niklas, 1992). On the other hand, the pectin into cell walls has a gel-like behavior that can enhance the extensibility of roots under tensile loads. Our data showed that D10 had almost 2-fold lower Young‘s module compared with C10 while at 45-days well-watered treatment displayed 2-fold lower Young‘s module than extreme drought treatment, showing the opposite behavior (Table 1). Collapsible and folding cell walls are produced by local mechanical stress when the succulent tissue water content decreases, but succulent cells can maintain higher elasticities during drought events through cell wall remodeling (i.e., elastic adjustment) (Fradera-Soler et al., 2022; Ahl et al., 2019) and this process could be related with our root biomechanical findings. However, the osmotic adjustment, turgor pressure, cell wall mechanics, and growth during extreme drought periods remains unknown and continue to be enigmatic for many plant species, especially desert succulents.

## 5. Conclusions

In this study we investigated the *Opuntia ficus-indica* root biomechanical, hydraulic, and anatomical changes under 10-, 30-, and 45-days extreme drought periods. We found several differences between well-watered and extreme drought treated plants. Abrupt reduction in *Lp*_*r*_ were discovered at 10-, 30, and 45-days reaching almost 87% reduction at 45-days of extreme drought treatment compared with well-watered plants. Also, several differences in root elasticity and stifness were observed between well-watered and drought treatments, suggesting that at 10-days of extreme drought treatment roots become stiffer and less elastic compared to well-watered plants. Interestingly, at 45-days of extreme drought treatment the opposite happens, where root become more elastic and less stiff than control. This behavior could be related to the loss of water content and a reduction of *C*_tlp_ of root cells, and probably by the cell wall material modification under extreme drought treatments. The results presented here highlight the need of including key biomechanical, hydraulic, and anatomical information on the current knowledge of succulent root physiological behavior under extreme drought treatments, especially for unstudied tissues. These results could be adapted to other succulent (e.g., Atacama Desert of Chile species), and agronomic (e.g., avocado, grapevine) species. The combination of the used techniques and approaches can be used to innovate in the root biophysics area and create new opportunities for a deeper understanding of fine roots under drought.

## 5. ACKNOWLEDGEMENTS

This study was funded by Agencia Nacional de Investigación y Desarrollo (ANID), within the Fondecyt grant N° 1220235. C.B-S received funding through an ANID doctoral scholarship Nº21221119. The authors kindly thank E. Ponce for his assistance during the experiments.

## 6. AUTHOR CONTRIBUTIONS

V.H.P., C.B.S., and I.F.C. designed the experiments. V.H.P. performed the hydraulic conductivity experiments. V.H.P. performed the tensile strength testing. C.B.S. performed the bright field light microscopy imaging. C.B.S., D.C.C., and I.F.C. analyzed data. C.B.S., and I.F.C wrote the initial draft of the article; V.H.P., D.C.C. and I.F.C. revised and edited the article. I.F.C. supervised the project.

## 7. CONFLICTS OF INTERESTS

The authors declare that this research was conducted in the absence of any commercial or financial relationships that could be constructed as a potential conflict of interest.

## Supplementary material

### Supplementary material 1

*Opuntia ficus-indica* propagation and maintenance. (A), *Opuntia* cladodes were collected and transplanted into 1.1 L plastic pots filled with coarse quartz sand. *Opuntia* cladodes were maintain approximately at 16 to 26ºC, 30% relative humidity and 500-600 µmol photons m^-2^ s^-1^, and 16h-light/8h-dark cycle. (B), Root proliferation from nodes under acclimatization process.

### Supplmentary material 2

Representative root scans showing the different heights (A-F) and inlets represent the widths. Differences in root diameter were found between well-watered and 45-days old extreme drought period plants. (F), Root shrinkage can reach to 22%.

### Supplementary material 3

Root tensile strength setup using a Ta.XT plusC texture analyzer. *Opuntia* root cortex failure and stele mechanical resistance until break under axial loading.

## Notes

### Competing Interest Statement

The authors have declared no competing interest.

